# Transcriptomic alterations including p53 pathway dysregulation prime DNMT3A mutant cells for transformation

**DOI:** 10.1101/2024.08.12.607467

**Authors:** Erin M Lawrence, Amali Cooray, Andrew J Kueh, Martin Pal, Lin Tai, Alex Garnham, Connie Li Wai Suen, Hannah Vanyai, Quentin Gouil, James Lancaster, Sylvie Callegari, Lauren Whelan, Elizabeth Lieschke, Annabella Thomas, Andreas Strasser, Yang Liao, Wei Shi, Andrew Wei, Marco J Herold

## Abstract

*DNMT3A* mutations are prevalent in haematologic malignancies. Our mouse model introduced the murine homologue (R878H) of the human ‘hotspot’ R882H mutation into the mouse *Dnmt3a* locus, resulting in globally reduced DNA methylation in all tissues. Mice with heterozygous R878H mutations developed γ-radiation induced thymic lymphoma more rapidly than control mice, suggesting a vulnerability to stress stimuli in *Dnmt3a^R878H/+^* cells. In competitive transplantations, *Dnmt3a^R878H/+^* Lin^-^Sca-1^+^Kit^+^ (LSK) cells had a competitive advantage over wt cells, indicating a self-renewal phenotype at the expense of differentiation. RNA-sequencing of *Dnmt3a^R878H/+^* LSKs exposed to low dose γ-radiation showed downregulation of the p53 pathway. Accordingly, reduced PUMA expression was observed by flow cytometry in the bone marrow of γ-irradiated *Dnmt3a^R878H/+^* mice due to altered p53 signalling. These findings provide new insights as to how *DNMT3A* mutations cause subtle changes in the transcriptome of LSK cells which contribute to their increased self-renewal and propensity for malignant transformation.

**SIGNIFICANCE:** Hotspot *DNMT3A* R882H mutations are overrepresented in leukaemia suggesting that they confer a susceptibility to disease if further mutations are acquired. To advance therapies for *DNMT3A* mutant disease, it is essential to understand the mechanisms by which this mutation primes cells for malignant transformation.

## INTRODUCTION

Mutations in epigenetic modifiers are common tumour initiating mutations in hematologic cancers. One of the most frequent mutations present in hematologic malignancies occurs in DNA methyltransferase 3 alpha (*DNMT3A*) [1–3]. While a mutation in *DNMT3A* alone is not sufficient to cause malignancy [4, 5], it is a potent driver of clonal haematopoiesis (CH) [6]. It is widely understood that *DNMT3A* is essential for maintaining the balance between hematopoietic stem cell (HSC) self-renewal and differentiation [7]. Genetic loss of *DNMT3A* causes an almost indefinite HSC self-renewal phenotype [7–9], which may explain its activity as a common driver of CH in humans. However, not all *DNMT3A* mutations are functionally equal. There is a considerable bias towards mutations at arginine 882 (*DNMT3A^R882^*) [4, 10], most commonly an R882H mutation, indicating that this mutation confers a possible competitive advantage to the cell. Indeed, *DNMT3A^R882H^* mutations are associated with poor patient outcomes in acute myeloid leukemia (AML), highlighting the increased pathogenicity of the *DNMT3A^R882H^* mutation in patients and a critical need for targeted therapies [11, 12].

The effect of *DNMT3A^R882^* in a pre-leukemic setting is not well understood. The murine homolog of the R882H mutation is R878H, and *Dnmt3a^R878H^* mouse models [13, 14] have previously been generated to examine the impact of this mutation in the initiation of CH and AML. To extend the work of others [15, 16] we used CRISPR/Cas9 gene editing to generate a *Dnmt3a^R878H/+^* mouse model. Unlike Cre inducible *Dnmt3a^R878H^* models, this allowed us to investigate the impact of *DNMT3A^R878H^* mutations with minimal intervention, avoiding the risk that Cre recombination itself could impact the phenotype. Using this model, we explored phenotypic and transcriptomic changes in pre-leukemic cells that caused the accumulation of clones in the blood, and investigated why these cells are at an increased risk of malignant transformation.

*DNMT3A* mutations may also have unrecognized non-canonical functions in HSCs. There is emerging literature associating p53 dysfunction with mutant *DNMT3A*, although it is unclear if this contributes to any disease phenotypes [17–19]. By exploring how changes in the epigenome and transcriptome may contribute to the initiation of malignancy in *DNMT3A* mutant cells, we identify disease mechanisms that are specific to *DNMT3A* mutant cells which will aid the development of therapies that will improve the survival of patients with *DNMT3A* mutant driven pathologies.

## RESULTS

### Generation of a *Dnmt3a^R878H/+^* mouse model using CRISPR/Cas9 gene editing technology

The *Dnmt3a^R878H/+^* mouse model was created using CRISPR/Cas9 gene editing technology **(Figure 1A).** Male *Dnmt3a^R878H/+^* mice were inter-crossed with female C57BL/6 mice to generate heterozygous *Dnmt3a^R878H/+^* mice. As previously reported [16], female *Dnmt3a^R878H/+^*mice had birthing difficulties and were not used for breeding.

**Figure 1.**
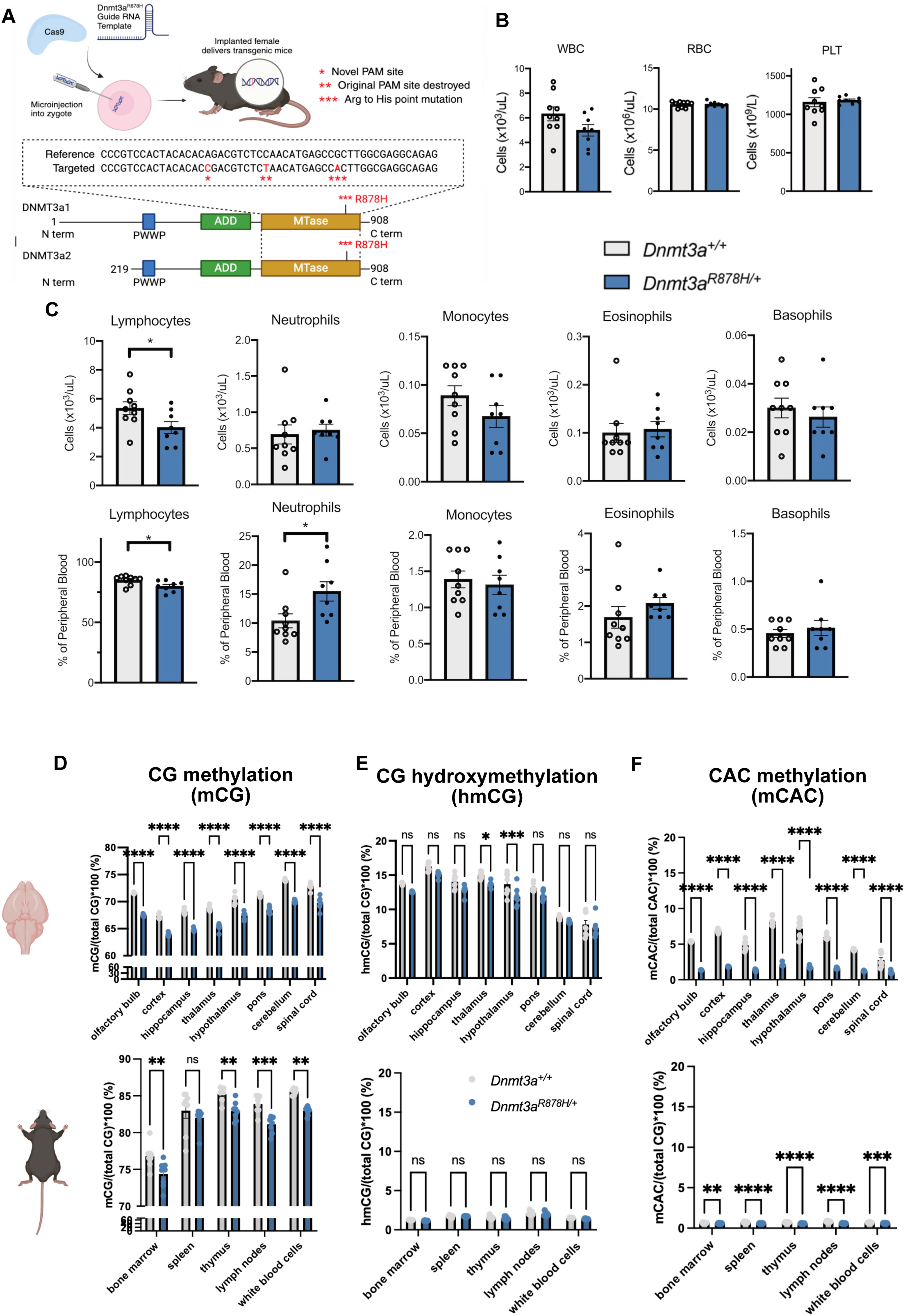
*Dnmt3a^R878H/+^* mice have reduced DNA methylation but normal hematopoietic cell populations. **A)** Schematic depicting the design of the *Dnmt3a^R878H^* mouse model. Three point mutations were introduced using CRISPR/Cas9 gene editing technology to create a novel PAM allowing the mutant allele to be specifically targeted (*), destroy the original PAM site to prevent Cas9 targeting the correct allele (**)_ and introduce the *Dnmt3a^R878H^* mutation into exon 23 of the *Dnmt3a* gene (***). **B)** White blood cell (WBC), red blood cell (RBC) and platelet (PLT) counts from *Dnmt3a^R878H/+^* (n=8) and WT (n=9) 8-10 week old mice, collected by retro-orbital bleeds. Cell counts determined by ADVIA analyzer at time of death. Students t-test, * p<0.05. **C)** Mature blood cells in *Dnmt3a^R878H/+^* and WT mice presented by both number and by percentage. Students t-test, with *= p<0.05. **D-F)** Different methylation readouts: mCG (D), hmCG (E), and mCAC (F), presented as a percentage of total CG in *Dnmt3a^R878H/+^* and WT brain regions and body tissues (n=8 WT, n=8 *Dnmt3a^R878H/+^*). %mCG is presented with a break in the y-axis to better visualise differences between genotypes. Data were statistically analysed using a 2-way ANOVA and Šídák’s multiple comparisons test, with *=p<0.0332, **=p<0.0021, ***=p<0.0002, ****=p<0.0001.

To determine the impact of *Dnmt3a^R878H/+^* on haematopoiesis, peripheral blood (PB) analysis was performed on *Dnmt3a^R878H/+^*mice and wild-type (WT) littermates at 8-10 weeks of age. There were no significant differences in white blood cell (WBC), red blood cell (RBC) or platelet counts **(Figure 1B)**. However, *Dnmt3a^R878H/+^* mice had fewer lymphocytes by both percentage and absolute number, and this coincided with a proportional increase in neutrophils by percentage, while other blood cell populations appeared unaffected by the mutation **(Figure 1C).** Further analysis of the blood and spleen by flow cytometry revealed no additional abnormalities in the *Dnmt3a^R878H/+^* mice **(Supplementary Figure 1A, 1B)**.

### *Dnmt3a^R878H/+^* mice exhibit reduced DNA methylation across several organs

The consequence of the *Dnmt3a^R878H/+^* mutation on global DNA methylation was assessed via low-coverage direct DNA sequencing [20]. The epigenetic process of DNA methylation involves the transfer of a methyl group to carbon 5 of a cytosine, forming 5-methylcytosine (5mC). The addition of a hydroxyl group to 5mC generates 5-hydroxy-methyl-cytosince (5hmC), which may be an intermediate stage in DNA demethylation that can contribute to regulation of gene expression [21, 22]. In mammals, CG methylation (mCG) which occurs at cytosines preceding guanines, also called CpG sites, is the most common type. Non-CpG methylation, such as in the methylated cytosine-adenosine (mCA) dinucleotide context, occurs less frequently, and primarily in the brain [23] and in pluripotent stem cells [24, 25]. Several lymphoid tissues, as well as the brain, which was further divided into regions, were collected from 8 *Dnmt3a^R878H/+^* mice and 8 WT littermates, the genomic DNA extracted, and an Oxford Nanopore Technologies’ nanopore sequencer was used to perform sequencing of extracted DNA from each tissue at low coverage (0.01-0.05X) to measure the global levels of cytosine modifications on autosomes. Apart from the spleen, every tissue measured had a consistent reduction of mCG in *Dnmt3a^R878H/+^* mice compared to WT littermates **(Figure 1D)**. The reduction in methylation was strongest in the brain regions, and more subtle in other tissues. This global hypomethylation phenotype is consistent with previous reports [8, 26, 27]. Hydroxymethylation of CpG sites (hmCG) was observed to be significantly reduced in the thalamus and hypothalamus of *Dnmt3a^R878H/+^* mice, while there was no significant reduction in any lymphoid tissues examined, which have much lower prevalence of hmCG compared to the brain [28] **(Figure 1E)**. Notably, CAC methylation (mCAC), a highly specific DNMT3A-catalysed modification [25], was substantially reduced in *Dnmt3a^R878H/+^* mice across all tissues tested **(Figure 1F)**.

### *Dnmt3a^R878H/+^* hematopoietic stem and progenitor cells are primed for γ-irradiation induced thymic T cell lymphomagenesis

A *DNMT3A* mutation may be among the first to arise in the progression from normal to malignant haematopoiesis [29], but the mechanism of action is incompletely understood. It has been suggested previously that mutant *DNMT3A* causes an impaired DNA damage response in pre-leukaemic cells [12]. To test this, we used a model of γ-radiation induced thymic lymphoma development (**Figure 2A)**. In this model, repeated low dose γ-irradiation was shown to promote leukemic transformation by acting on early haematopoietic stem and progenitor cells (HSPCs) in the bone marrow [30]. These nascent leukemic cells migrate from the bone marrow to the thymus where their expansion drives T cell lymphoma development.

**Figure 2:**
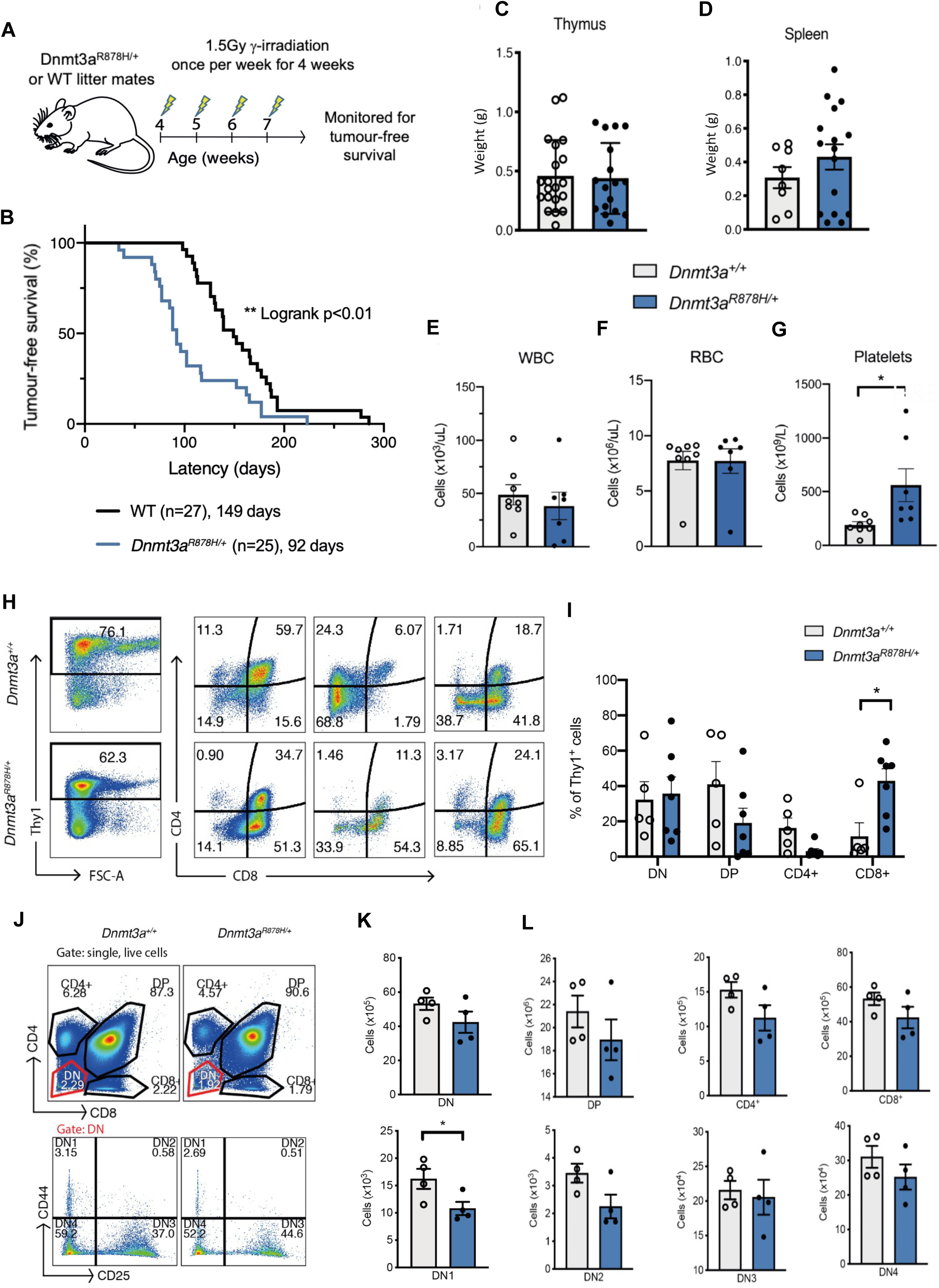
*Dnmt3a^R878H/+^* hematopoietic stem and progenitor cells are primed for γ-radiation induced thymic lymphoma. **A)** Schematic indicating *Dnmt3a^R878H/+^* mice and WT control littermates exposed to four doses of 1.5 Gy γ-radiation, once a week, starting the age of 4 weeks with the aim of producing γ-irradiation-induced thymic lymphoma. **B)** Kaplan-Meier survival curve comparing tumor-free survival of *Dnmt3a^R878H/+^*mice compared to WT control littermates. Data were statistically analysed using a log rank (Mantel-Cox) statistical test, with **=p<0.01. **C)** Thymus and **D)** spleen weights respectively, of *Dnmt3a^R878H/+^* (n=16) and WT (n=20) mice euthanised at the ethical endpoint after developing thymic lymphoma. Data were statistically analysed using a Student’s t-test. **E)** White blood cells (WBC), **F)** red blood cells (RBC), and **G)** platelet (PLT) counts from *Dnmt3a^R878H/+^* (n=8) and WT (n=7) thymic lymphoma-burdened mice at time of death. Data were statistically analysed using a Students t-test, *=p<0.05. **H)** Flow cytometric plots and **I)** graphical representations of the distribution of CD4 and CD8 expressing thymic lymphoma cells. *Dnmt3a^R878H/+^-*derived lymphomas were significantly more likely to express CD8 compared to WT-derived tumor cells. This was accompanied by a modest reduction in double-positive CD4^+^CD8^+^ lymphoma cells and single positive CD4^+^ lymphoma cells although this did not reach statistical significance. Data were statistically analysed using Multiple t-tests, *=p<0.05. **J)** Flow cytometry gating of CD4 and CD8 single-positive T cells, double-positive (DP) T cells and double-negative (DN) T cells from thymic lymphomas (top) with the DN T cells further divided into their differentiation stages (bottom; DN1 (CD25^-^CD44^+^), DN2 (CD25^-^CD44^+^), DN3 (CD25^+^CD44^-^) and DN4 (CD25^-^CD44^-^)). **K)** Graphical representations of the numbers of DN thymic lymphoma T cells, and those in DN1, from *Dnmt3a^R878H/+^* (n=4) and WT (n=4) mice. Data were statistically analysed using Student’s t-tests, *=p<0.05. **L)** Graphical representation of the more mature T cell numbers from thymic lymphomas, as well as those in DN2-4, from *Dnmt3a^R878H/+^* (n=4) and WT (n=4) mice Data were statistically analysed using Student’s t-tests.

*We observed that Dnmt3a^R878H/+^* mice developed thymic T cell lymphoma significantly faster than their WT littermates **(Figure 2B)**. Importantly, it appeared that the *Dnmt3a^R878H/+^* mice had a similar tumour burden to WT mice at death, as indicated by non-significantly different weights of their abnormally enlarged thymus and spleen **(Figures 2C, 2D)**. PB analysis at the point of sacrifice also revealed no significant difference in WBC and RBC counts between *Dnmt3a^R878H/+^*and WT mice **(Figures 2E, 2F)**. However, *Dnmt3a^R878H/+^* mice had significantly higher platelet counts (∼2-fold) compared to WT controls **(Figure 2G)**. No significant differences were found in other blood cell types **(Supplementary Figures 1C-H)**. Flow cytometric analyses of spleens revealed no significant differences in their cell composition between *Dnmt3a^R878H/+^* and WT controls **(Supplementary Figure 1I).**

To identify possible phenotypic differences between *Dnmt3a^R878H/+^-* and WT-derived tumors, flow cytometric analysis was performed on the thymus of the tumor burdened mice. While there was no difference in TCRβ and B220 expression **(Supplementary Figures 1J & K)**, *Dnmt3a^R878H/+^*derived lymphomas were significantly more likely to express CD8 compared to WT lymphomas **(Figures 2H, 2I)**. This data suggests that DNMT3A may act as a CD8 T cell lymphoma suppressor, and/or regulate T cell differentiation, the latter in line with previous reports [31, 32]. The CD8^+^ *Dnmt3a^R878H/+^*lymphomas may represent malignant counterparts of immature T lymphoid cells in the thymus that express CD8 prior to becoming CD4/CD8 double positive (DP).

This may indicate that mutant DNMT3A causes a defect in T cell differentiation in the thymus. To examine this, the subcellular composition of the thymus was analysed in healthy (i.e. non-irradiated) *Dnmt3a^R878H/+^*and control WT mice. There were significantly fewer double-negative differentiation stage 1 (DN1) thymocytes in *Dnmt3a^R878H/+^* mice compared to WT controls **(Figures 2J, 2K)**. *Dnmt3a^R878H/+^* thymocytes maintained a trend towards reduced cell numbers throughout the later DN stages, as well as the more mature CD4 and CD8 single positive cell subsets, however these reductions did not reach statistical significance **(Figure 2L)**. Collectively, this data may indicate that mutant DNMT3A may have a subtle influence on the differentiation of early progenitor populations in the thymus, and possibly the bone marrow.

### *Serial competitive bone marrow transplantations demonstrate Dnmt3a^R878H/+^*hematopoietic stem and progenitor cells have a competitive advantage over WT cells

Flow cytometric analysis of *Dnmt3a^R878H/+^* and WT littermate bone marrow HSC compartments (**Supplementary Figure 2A**) revealed no significant differences in the numbers of granulocyte myeloid (GMP), common myeloid (CMP) or megakaryocyte erythroid (MEP) progenitors, although there was a trend towards slightly higher numbers of each of these progenitor populations in the *Dnmt3a^R878H/+^* mice (**Figure 3A**). No significant differences were found between the numbers of long term- or short term-HSCs (LT-/ST-HSCs) between *Dnmt3a^R878H/+^* and WT mice (**Figure 3B**). This again indicates any effect of the *Dnmt3a^R878H/+^* point mutation on HSCs is quite subtle, as has been reported in previous literature [33].

**Figure 3:**
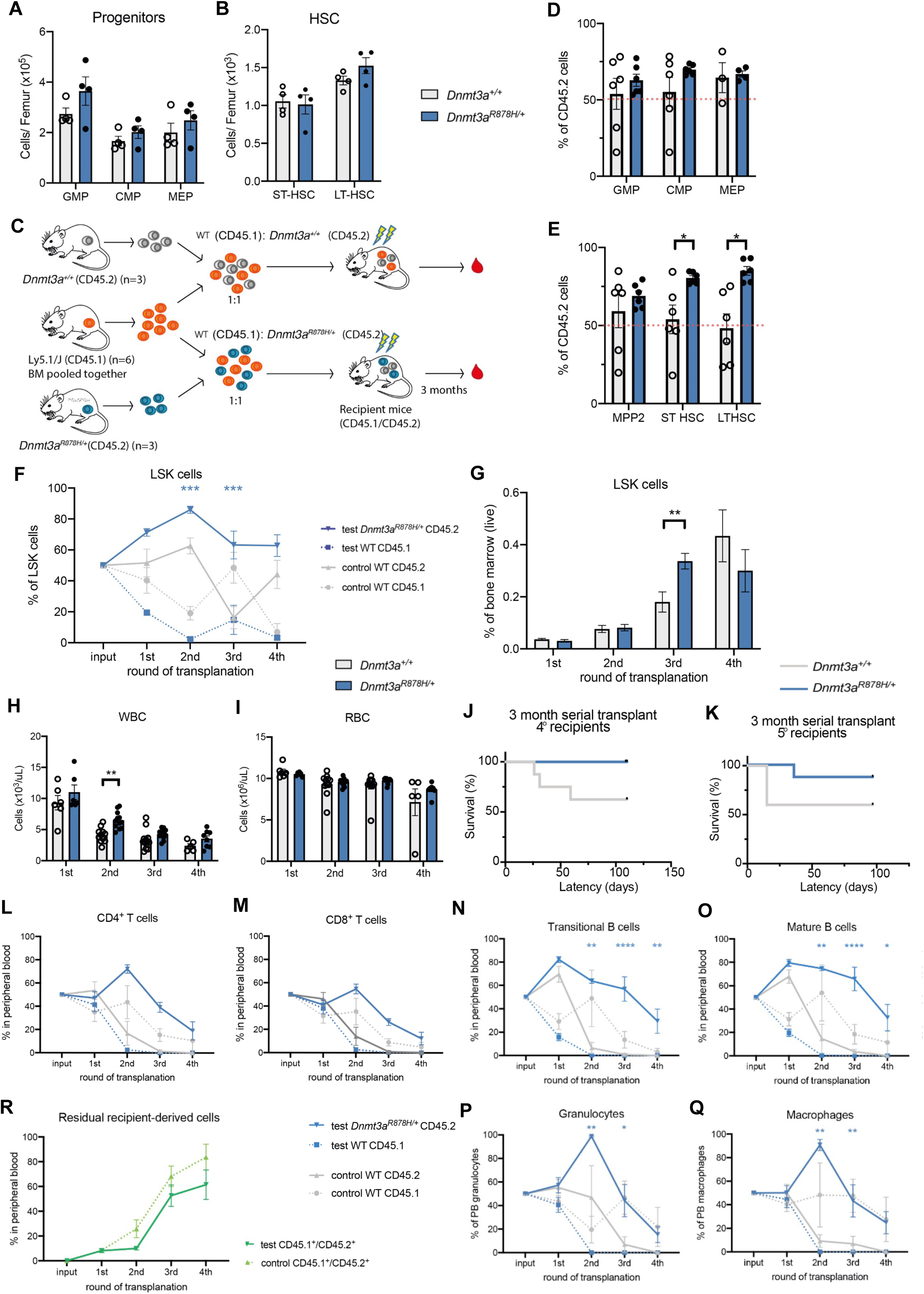
*Dnmt3a^R878H/+^*hematopoietic stem and progenitor cells have a competitive advantage over WT cells as shown by serial competitive bone marrow transplantation. **A)** Progenitor cells and **B)** hematopoietic stem cells numbers per femur from 8-12 week old *Dnmt3a^R878H/+^* (n=4) and WT control (n=4) mice. Data were statistically analysed using Student’s t-tests, *=p<0.05. **C)** Schema depicting the experimental setup of the competitive bone marrow transplantation experiments. **D), E)** The proportions of CD45.2 HSPCs (granulocyte-myeloid progenitors (GMP), common myeloid progenitors (CMP), megakaryocyte-erythroid progenitors (MEP) (D), and multipotent progenitor cells (MPP), long-term hematopoietic stem cells (LT-HSC), and short-term haematopoietic stem cells (ST-HSC) (E)) from *Dnmt3a^R878H/+^* (n=6) and WT control (n=6) recipient mice three months after the original haematopoietic reconstitution transplant. **F)** The proportions of donor CD45.2 LSK cells in *Dnmt3a^R878H/+^*mice or WT recipient mice 3 months after each round of transplantation **G)** Graph showing the proportions of donor LSK cells found in mice that received the combined WT (CD45.1)/ *Dnmt3a^R878H/+^* (CD45.2) bone marrow 3 months after each round of transplantation. **H)** WBC and **I)** RBC number from mice 3 months after each serial transplantation with WT:WT or WT:*Dnmt3a^R878H/+^* bone marrow cell mixtures. Data were statistically analysed using Student’s t-tests, **=p<0.01. **J)** Kaplan-Meier survival curve of fourth generation (4°) recipients of serially transplanted bone marrow. Data were statistically analysed using a log-rank (Mantel-Cox) test. **K)** Kaplan-Meier survival curve of fifth generation (5°) recipients of serially transplanted bone marrow. Data were statistically analysed using a log-rank (Mantel-Cox) test.. Blood cell proportions determined by flow cytometry for **L)** CD4+ T cells, **M)** CD8+ T cells, **N)** transitional B Cells, **O)** mature B Cells, **P)** granulocytes, and **Q)** macrophages, all from mice (n=8 per group) 3 months after each serial transplantation. Data were statistically analysed using two-way ANOVAs with Šídák’s multiple comparisons test. **R)** Proportions of recipient-derived cells over time in the WT:*Dnmt3a^R878H/+^* reconstituted mice compared with the WT:*Dnmt3a^+/+^* reconstituted control mice.

Previous studies have established that *DNMT3A*-KO HSPCs have a propensity to self-renew at the expense of differentiation [7, 8, 29, 34]. To determine whether this held true for *Dnmt3a^R878H/+^* HSPCs, we conducted competitive bone marrow transplantation experiments **(Figure 3C)**. CD45.2 *Dnmt3a^R878H/+^* or WT bone marrow were combined in a 1:1 ratio with WT CD45.1 cells, and used to reconstitute the haematopoietic system of lethally irradiated CD45.1^+^CD45.2^+^ recipient mice. After 3 months of recovery, the proportions of donor progenitor cells was determined based on CD45 isoform surface marker expression. While no significant difference in the proportions of *Dnmt3a^R878H/+^* derived cells was observed in GMP, CMP and MEP populations, (**Figure 3D**), *Dnmt3a^R878H/+^* LT- and ST-HSCs were strongly enriched in the bone marrow of recipient mice compared to their control WT counterparts (**Figure 3E**). This is in line with previous reports of *Dnmt3a^R878H/+^* driven clonal expansion [15, 35] and supports the notion that mutant *DNMT3A* plays an important role in the balance between self-renewal and differentiation of stem cells.

To test whether the advantage of *Dnmt3a^R878H/+^* HSCs could be sustained, we isolated the bone marrow from the reconstituted recipient mice at 3 months and repeated the transplantation as previously described, into lethally irradiated secondary recipient mice. This transplantation process was repeated until the recipient mice could no longer be rescued from lethal irradiation. By examining the LSK (Lin^−^ Sca-1^+^ c-Kit^+^) compartment in the bone marrow after each round of transplantation it was found that *Dnmt3a^R878H/+^* LSK cells had a sustained competitive advantage over at least 4 rounds of serial transplantation (**Figure 3F**). However, the increased proportion of *Dnmt3a^R878H/+^* LSK cells did not equate to an uncontrolled proliferation of cells, as the LSK cells still made up a similar proportion of the total bone marrow when compared to WT control cells (**Figure 3G**), the exception being in the third transplant, although this returned to equivalent percentages in the fourth round.

Furthermore, despite *Dnmt3a^R878H/+^* LSK cells outcompeting WT LSK cells in in the bone marrow of reconstituted mice, the overall blood composition remained similar between the WT:WT control and the WT:*Dnmt3a^R878H/+^* mixed bone marrow recipients. Mice receiving the WT:*Dnmt3a^R878H/+^* bone marrow mix had significantly higher WBC counts after the second transplant, however no significant differences were otherwise detected (**Figures 3H, 3I**). By the 4^th^ and 5^th^ rounds of serial transplantation, HSC exhaustion began to cause lethal hematopoietic ablation. Importantly, the mice that had received the WT:*Dnmt3a^R878H/+^* bone marrow were more protected from this exhaustion as shown by the first reconstitution failure occurring only in the 5^th^ round of transplantation (**Figures 3J, 3K**).

When assessing the contribution of donor cells to more mature cell subtypes in the PB, *Dnmt3a^R878H/+^* cells were more likely to make up a larger proportion of both CD4 and CD8 T cells at all rounds of transplantation measured. However, this did not reach statistical significance, perhaps due to a higher number of γ-radiation resistant host-derived T cells that remained in these mice (**Figures 3L, 3M**). Furthermore, there was a strong and sustained competitive advantage of both transitional and mature B cells as well as granulocytes and macrophages from the *Dnmt3a^R878H/+^* donors (**Figures 3N-Q**), indicating that *Dnmt3a^R878H/+^* mutant cells do not have a lymphoid or myeloid bias under these conditions. Across all mature cell types examine, it appeared as though the *Dnmt3a^R878H/+^* cells lost their competitive advantage after multiple rounds of transplantation as seen by their reduction in proportion in the PB (**Figure 3L-Q**). However, this coincided with an increase in the contribution of residual recipient derived (CD45.1^+^/CD45.2^+^) cells (**Figure 3R**). These cells likely had escaped γ-radiation and reflect ‘fitter’ cells compared with the serially transplanted mixed bone marrow cells.

### *Dnmt3a^R878H/+^* myeloid progenitor cells have a subtle survival advantage upon γ-irradiation

The serial competitive transplantations demonstrated that although *Dnmt3a^R878H/+^* cells had a distinct survival advantage, no myeloid or lymphoid bias was seen in steady state conditions. To explore how *Dnmt3a^R878H/+^* could be interfering with HSCs, we analysed HSC survival after exposure to γ-irradiation. Bone marrow was harvested from 8-12 week old *Dnmt3a^R878H/+^* and WT littermates, cells transferred into cell culture medium [36], exposed *ex vivo* to γ-irradiation (1.5 Gy) or left untreated. Cell viability was then assessed by flow cytometry over 72 h with the HSC gating strategy [37] shown in **Supplementary Figure 2B**. *Dnmt3a^R878H/+^* HSCs initially trended towards slightly better survival compared to their WT cells, and this advantage appeared subtly more pronounced in the γ-irradiated cells (**Figure 4A**). *Dnmt3a^R878H/+^* LSK cells showed a trend towards better survival compared to their WT counterparts at 12 h, although no survival advantage was observed at later timepoints (**Figure 4B**). No statistically significant differences were found between the HSC compartments when when looking at total cell numbers across all timepoints. Non irradiated *Dnmt3a^R878H/+^* ST- and LT-HSCs trended toward better survival than their WT counterparts, though this did not reach statistical significance **(Supplementary Figures 3B-D)**. The *Dnmt3a^R878H/+^* myeloid progenitor cells (MPP3) were observed to be slightly more robust following γ-irradiation compared to their WT counterparts although this was not also statistically significant **(Supplementary Figure 3A).** These data again support *Dnmt3a^R878H/+^* stem/progenitor cells having a subtle competitive advantage, and a similarly subtle survival advantage in myeloid progenitors upon DNA damage.

**Figure 4:**
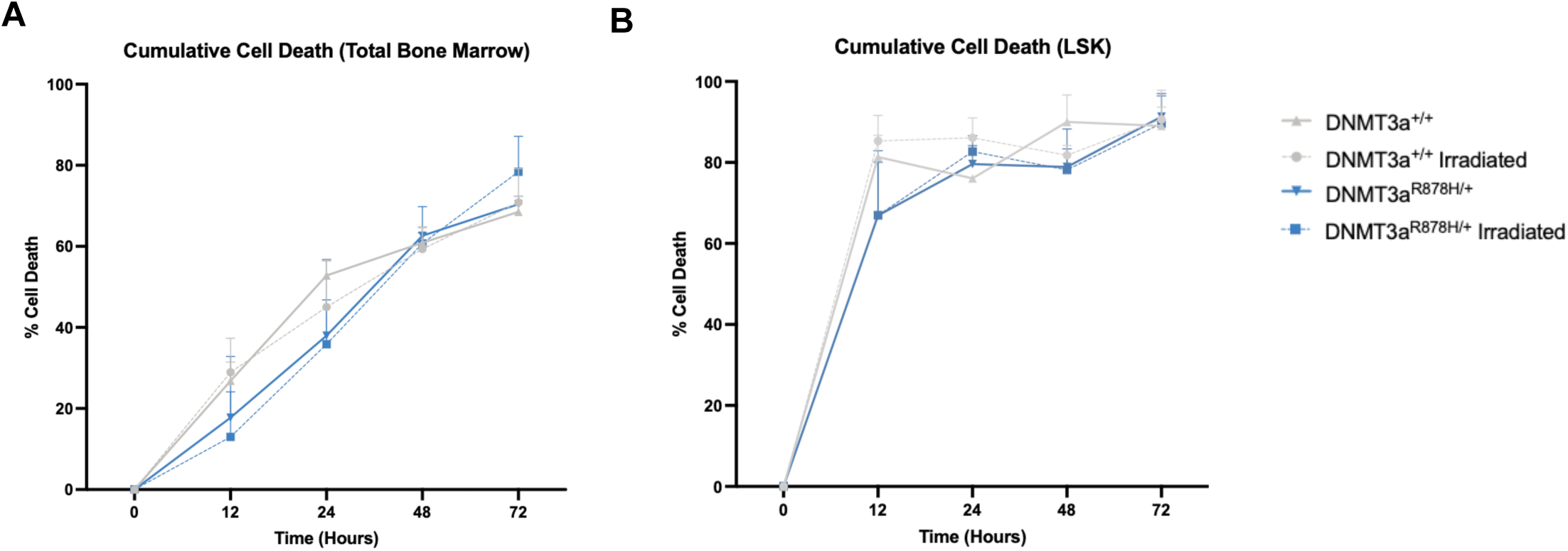
*Dnmt3a^R878H/+^* cells have a subtle survival advantage upon γ-irradiation. Representations of cumulative cell death from flow cytometric analyses of **A)** total cells and **B)** LSK cells from *Dnmt3a^R878H/+^* and WT bone marrow, following 1.5Gy γ-irradiation or gnon-treated controls. Percentage of cell death is measured relative to time point 0. Data were statistically analysed using two-way ANOVAs with Šídák’s multiple comparisons test. For each group n=3 per 3 independent experiments.

### Transcriptomic profiling of γ-irradiated pre-leukemic *Dnmt3a^R878H/+^* LSK cells reveals subtle downregulation of p53 signalling pathway genes

We have demonstrated that *Dnmt3a^R878H/+^* mice succumb more rapidly to γ-irradiation induced thymic lymphoma. We hypothesized this may be due to an abnormal or incomplete stress response, and therefore explored the transcriptional changes taking place in response to γ-irradiation in *Dnmt3a^R878H/+^* and WT LSK cells via RNA-seq. *Dnmt3a^R878H/+^* and WT littermates were either left untreated or exposed to low dose, whole body γ-irradiation (2 Gy), and then allowed to recover for 2 hours before bone marrow was harvested and LSK cells were isolated by FACS. The RNA was then extracted from LSK cells and bulk RNA-seq was performed **(Figure 5A)**. When comparing transcriptional changes between LSK cells from γ-irradiated *Dnmt3a^R878H/+^* LSK cells with their γ-irradiated WT counterparts, 335 genes were observed to be differentially expressed (DE), with 105 downregulated and 230 upregulated DE genes identified (**Figure 5B, 5C)**.

**Figure 5:**
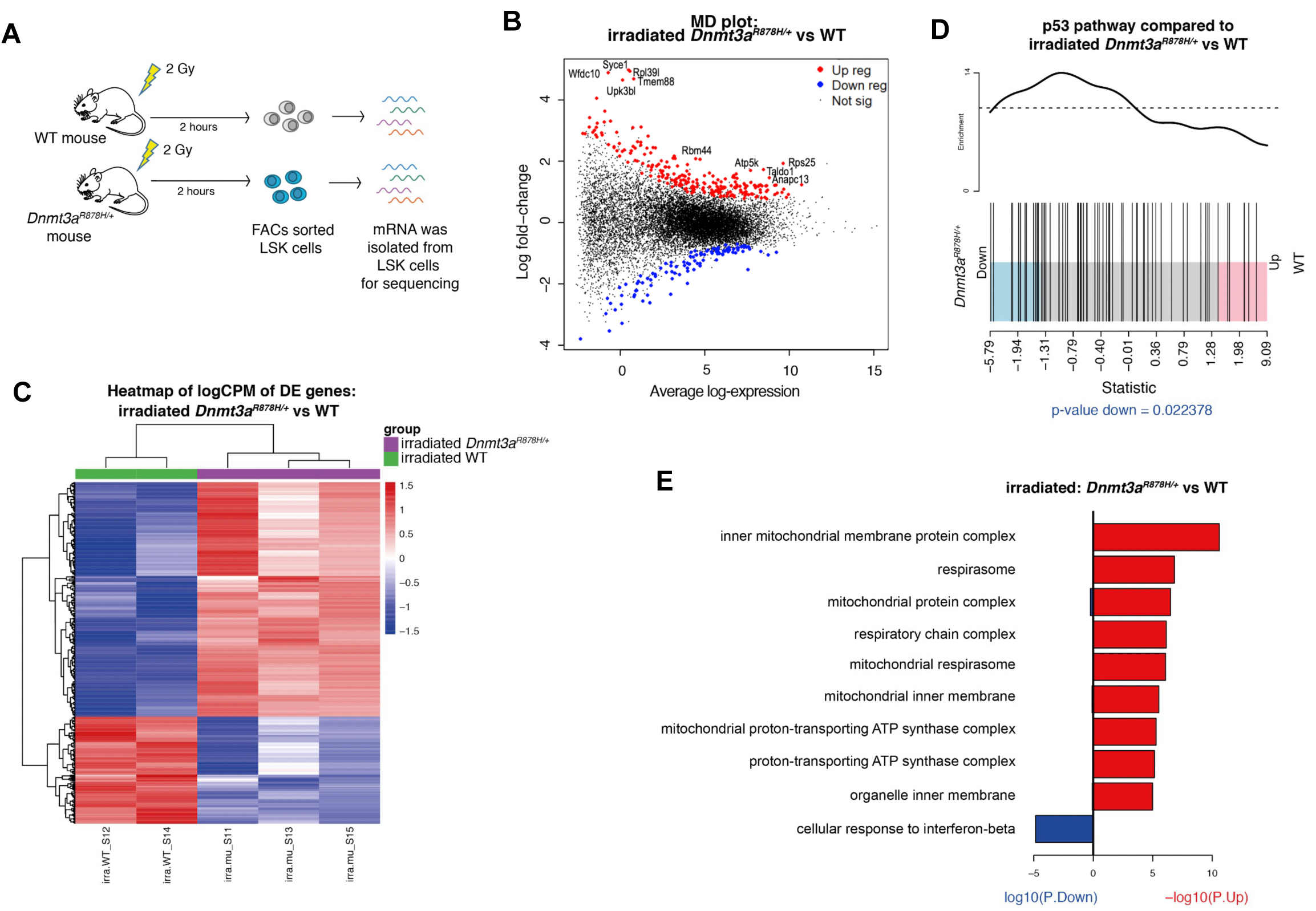
RNA sequencing of y-irradiated *Dnmt3a^R878H/+^* LSK cells reveals transcriptomic alterations including p53 pathway dysregulation. **A)** Schematic depicting the experimental protocol where *Dnmt3a^R878H/+^* and WT control littermates were exposed to a single dose of whole-body γ-radiation (2 Gy). The mice were allowed to recover for 2 hours before LSK cells were isolated from the bone marrow for RNAseq. **B)** MD plot depicting gene expression differences between γ-irradiated *Dnmt3a^R878H/+^* LSK cells *vs* γ-irradiated WT control LSK cells where genes in red are significantly increased and genes in blue are significantly decreased in *Dnmt3a^R878H/+^* LSK cells. **C)** Heatmap depicting differentially expressed genes between γ-irradiated *Dnmt3a^R878H/+^* LSK cells *vs* γ-irradiated WT control LSK cells. **D)** Barcode plot comparing p53 pathway gene expression between γ-irradiated Dnmt3a^R878H/+^ LSK cells and γ-irradiated WT LSK cells. **I)** The top 10 significantly enriched GO terms associated with differentially expressed genes in γ-irradiated *Dnmt3a^R878H/+^* LSK cells.

As expected, the p53 signalling pathway was significantly enriched in both *Dnmt3a^R878H/+^* and WT LSK cells upon γ-irradiation **(Supplementary Figures 4E, 4F)**. However, when compared to WT LSK cells, the p53 signalling pathway was significantly weaker induced in *Dnmt3a^R878H/+^* LSK cells, suggesting a blunted p53 response to DNA damage **(Figure 5D)**. GO pathway analysis further revealed a significant upregulation of gene transcriptional networks related to oxidative phosphorylation and various mitochondrial complex formation pathways in γ-irradiated *Dnmt3a^R878H/+^* LSK cells compared to their WT counterparts **(Figure 5E)**. However, Seahorse mito stress assays revealed no significant difference between the maximal oxygen consumption rate (OCR) of *Dnmt3a^R878H/+^* myeloid or B cells compared to their WT counterparts **(Supplementary Figures 5A-D).**

### Reduced PUMA induction in irradiated *Dnmt3a^R878H/+^* cells suggests an impaired p53 response

To further explore the weaker p53 pathway signalling observed in transcriptomic analysis of γ-irradiated *Dnmt3a^R878H/+^* LSK cells, we utilized a *Puma-tdTomato* reporter mouse [38] where tdTomato has been knocked into the *Puma/Bbc3* gene locus. *Puma/Bbc3* is a direct transcriptional target of p53 which is critical for p53 induced apoptosis [39]. Thus, *Puma* reporter expression was used as a measure of p53 transcriptional activity. Male *Dnmt3a^R878H/+^* mice were crossed with female *Puma*-*tdTomato^KI/KI^* reporter mice, and at 8-12 weeks, *Dnmt3a^R878H/+^/Puma-tdTomato^KI/+^* and *Dnmt3a^+/+^/Puma-tdTomato^KI/+^* littermates were either exposed to γ-irradiation (5Gy) or left untreated. 8 hours post-treatment, the spleen, thymus and bone marrow cells of the mice were collected, and tdTomato expression was measured by flow cytometry to observe *Puma* induction and p53 activity **(Figure 6A)**. As expected, γ-irradiation led to *Puma* reporter induction in all tissues examined in both *Dnmt3a^R878H/+^* and WT mice **(Figures 6B-D)**. The WT spleen and bone marrow cells showed significantly higher *Puma* reporter induction following γ-irradiation compared to their *Dnmt3a^R878H/+^* counterparts **(Figure 6E, 6F)**, while no significant difference was observed in the thymus **(Supplementary Figure 6A).** This supports the notion that *Dnmt3a^R878H/+^* LSK cells have blunted p53 activity, which may afford them a subtle survival advantage when sustaining DNA damage and this may also prime these cells for clonal expansion.

**Figure 6:**
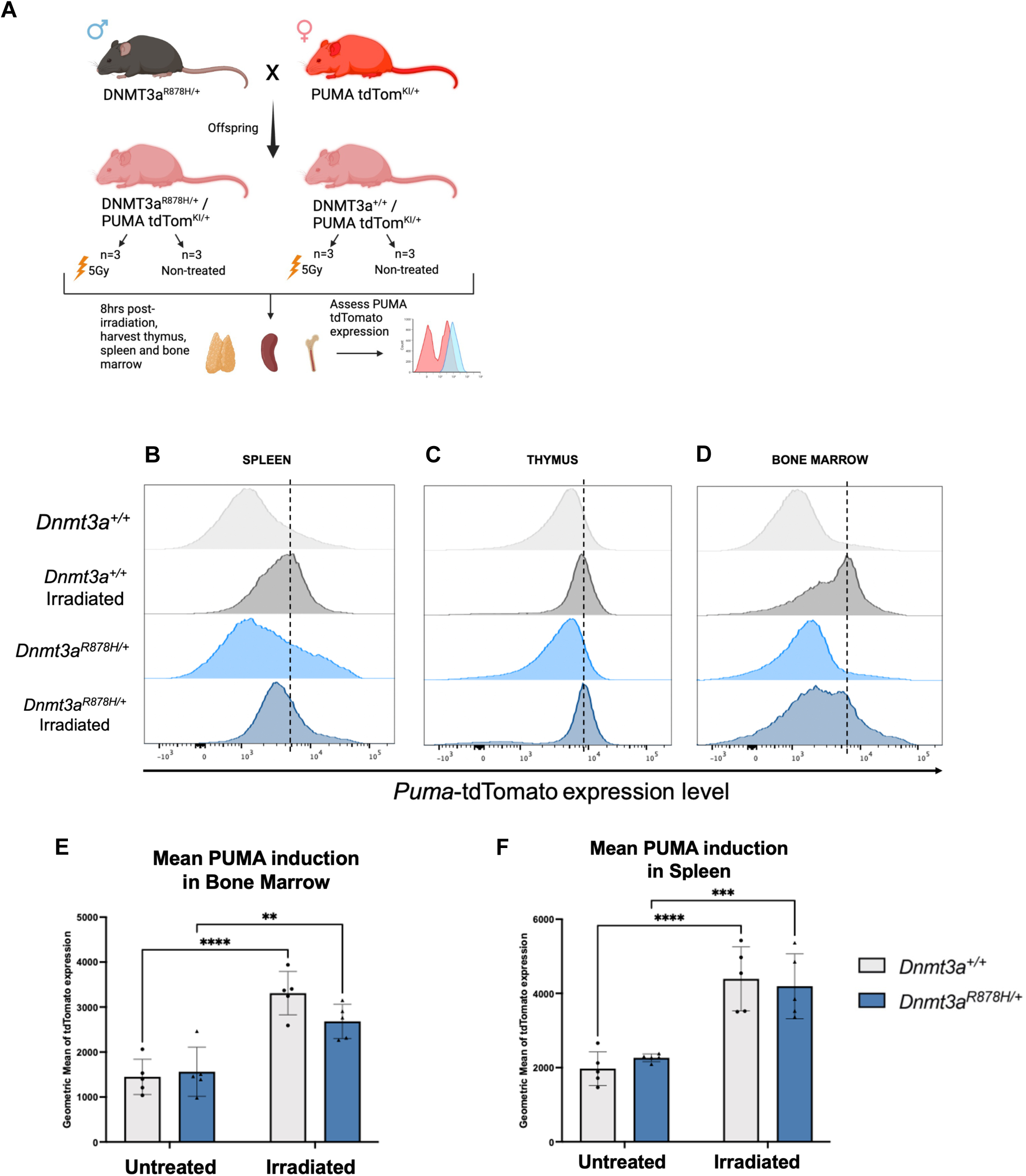
Reduced Puma-tdTomato reporter induction in γ-irradiated *Dnmt3a^R878H/+^* spleen and bone marrow cells is consistent with an impaired p53 response. **A)** Schematic for *Dnmt3a^R878H/+^* and *Puma-tdTomato* mating and experimental program. *Dnmt3a^R878H/+^; Puma-tdTomato^KI/+^* (n=6) and *Dnmt3a^+/+^; Puma-tdTomato^KI/+^* (n=6) offspring were either treated with 5Gy γ-irradiation (n=3 per genotype) or left untreated (n=3 per genotype). The thymus, spleen, and bone marrow from each mouse were then harvested and analysed for tdTomato expression by flow cytometry. **B-D)** Representative histograms showing tdTomato expression (reporting on *Puma* mRNA) in spleen (B), thymus (C) and bone marrow (D) cells in the γ-irradiated and non-irradiated groups. **E), F),** Graphical representations of geometric means of tdTomato expression in the bone marrow € and spleen cells (F) (n=5 per condition). Data were statistically analysed using two-way ANOVAs with Šídák’s multiple comparisons test, **=p<0.01, ***=p<0.001, ****=p<0.0001.

## DISCUSSION

There is a growing body of evidence for the importance of DNMT3A in haematopoiesis, showing that impairment of its function can initiate CH and leukemogenesis [3, 5, 29, 40]. In this study, we generated a mouse model that replicates the most common human *DNMT3A* mutation, the R882H substitution, by introducing the homologous murine mutation, R878H, into the *Dnmt3a* locus using CRISPR/Cas9 gene editing technology. An advantage of this model is that *Dnmt3a^R878H/+^* cells can be studied without interference from CRE mediated recombination. This strategy allowed us to study the impact of *Dnmt3a^R878H/+^* on haematopoiesis in intact mice and allowed us to isolate and study cell populations without the need for Cre mediated induction of the mutation. The data presented here support and extend findings from previous studies where expression of *Dnmt3a^R878H^* was studied in the context of normal and malignant haematopoiesis [13]. *DNMT3A* R882 mutations are now widely reported to be among the most common driver mutations in patients with AML [11, 41, 42]. However, as genomic sequencing becomes a mainstay in cancer diagnosis and research, mutant *DNMT3A* has also been found to occur in a range of lymphoid malignancies particularly among patients with adult T-ALL [43, 44].

A fast and efficient low coverage DNA sequencing approach revealed that our *Dnmt3a^R878H/+^* mice have reduced methylation across all tissues examined. This hypomethylation phenotype is especially pronounced in the brain, and more subdued in lymphoid organs examined. It has been shown that loss or alteration of *Dnmt3a* combined with a secondary cooperating lesion can induce leukemia in mice [45–47], but precisely how mutant DNMT3A contributes to the initiation of leukemia is still incompletely defined. There is also not yet a consensus in the field as to how the *DNMT3A* R882 hotspot mutation differs from deletion of the *DNMT3A* gene. Some groups proposed that mutant DNMTA R882 acts in a dominant negative fashion [48, 49], whereas others suggested that the hotspot mutation may alter the flanking sequence preference of *DNMT3A* [50, 51]. Realistically, it could be a contribution of both, alongside other features that have yet to be defined. It has been suggested that *DNMT3A* mutations can interfere with the ability of cells to sense and repair DNA damage, leading to increased mutagenesis [12].

We were able to test this hypothesis using our CRISPR engineered *Dnmt3a^R878H^*mice, as we could directly compare the effects of γ-irradiation-induced DNA damage *in vivo*. In line with previous studies [52], we demonstrated that *Dnmt3a^R878H^* had minimal effect on haematopoiesis at steady state. However, we observed a marked acceleration in γ-radiation induced thymic lymphoma development in *Dnmt3a^R878H/+^* mice compared to WT littermates. We know from previous work that the cell of origin of thymic lymphoma arises from a bone marrow derived progenitor cell [30]. This indicates that the accelerated lymphoma development in the *Dnmt3a^R878H/+^* animals likely is due to interference with the stress response in bone marrow HSPCs. Indeed, the work of others has shown that *Dnmt3a^-/-^* LSK cells have a pre-leukemic phenotype [29], and previous work on other murine models of *Dnmt3a^R878H/+^* LSK cells have also demonstrated pre-leukaemic traits in these cells [13]. Collectively, these studies suggest that mutant DNMT3A could drive a proliferative phenotype at the expense of differentiation. Therefore, we assessed whether we could also observe this phenotype in our CRISPR generated *Dnmt3a^R878H/+^* mutant mice.

Interestingly, we did not find any accumulation of *Dnmt3a^R878H/+^* HSPCs in the bone marrow of 8-week-old mice, indicating that any proliferative advantage conferred by mutant *DNMT3A* is subtle. To examine *Dnmt3a^R878H/+^* stem cell fitness we performed serial competitive bone marrow transplant experiments, which revealed that *Dnmt3a^R878H/+^* stem cells accumulate in the bone marrow, while any changes in mature blood cell proportions are only detectable 3 months after these transplantations. This finding was in support of previous research that revealed *Dnmt3a^R878H/+^* LSKs can lead to an increases abundance of *Dnmt3a^R878H/+^* mature leukocytes in PB [4]. Similarly, we observed an increase in the proportion of *Dnmt3a^R878H/+^* PB cells and showed that *Dnmt3a^R878H/+^* cells possess a survival advantage over their WT counterparts in the myeloid lineage. Our results further suggest that upon DNA damage, *Dnmt3a^R878H/+^* MPP3s are perhaps slightly more robust than their WT counterparts, and although the result was not statistically significant, it may suggest that these cells are better able to proliferate to compensate for the cell death seen in other stem cell as well as the mature compartments. As these cells proliferate, the acquisition of additional mutations over time could contribute to haematopoietic tumorigenesis.

To explore how this proliferative advantage of *Dnmt3a^R878H/+^* HSPCs could cause the increased propensity for leukemia development in the γ-radiation-induced thymic lymphoma model, we turned to transcriptomics. RNAseq revealed weakened activation of the p53 pathway in *Dnmt3a^R878H/+^* HSPCs following γ-irradiation. A *Puma-tdTomato* reporter mouse model was further used to confirm this impaired p53 transcriptional response, as reporter expression was significantly blunted following γ-irradiation in *Dnmt3a^R878H/+^* bone marrow and spleens compared to these tissues from WT mice.

Collectively, our findings show that *Dnmt3a^R878H/+^* drives altered gene expression patterns in LSK cells that have sustained DNA damage, providing a proliferative and survival advantage over WT cells. We propose that this advantage is driven at least in part by the attenuation of p53 activity in response to stress stimuli. The weakened p53 response may contribute to neoplasticity in mutant cells. Perhaps, boosting the p53 pathway may be a therapeutic target in CH and leukemias driven by mutant DNMT3A.

## CONCLUSION

To expand on current knowledge, we generated a *Dnmt3a^R878H/+^* mouse model to investigate the effect of the hotspot mutant DNMT3A in a pre-leukaemic setting. Our studies reveal that *Dnmt3a^R878H/+^* causes a broad hypo-methylation phenotype and drives the expansion of a pre-leukemic HSPC pool over time, such as during successive bone marrow transplantation. The *Dnmt3a^R878H/+^* cells display evidence of cancer-associated changes in p53 pathway gene expression consistent with pre-leukemic changes. Further exploration of the impact of this DNMT3A mutation on cell death and DNA damage repair may identify druggable targets. This may have implications for the prevention of *Dnmt3a^R878H/+^* driven CH and leukemias including for the treatment of minimal residual disease.

## MATERIALS AND METHODS

### Animals

The use of animals was approved by the Walter and Eliza Hall Institute of Medical Research (WEHI) animal ethics committee. The *Dnmt3a^R878H/+^***^ ^**mice and *Puma-tdTomato* reporter mice [38] were generated by the MAGEC laboratory at the Walter and Eliza Hall Institute (WEHI) as previously described by Kueh et al [53]. These mice were generated and maintained on a C57BL/6 background. For the *Dnmt3a^R878H/+^***^ ^**mice, Cas9 mRNA, a *Dnmt3a*-targeting single guide RNA (sgRNA) (5’-CCTCGCCAAGCGGCTCATGT-3’), and donor reference oligo (5’-ccgcactcactcccttccctgccttcctcccacagGGTGTTTGGCTTCCCCGTCCACTACACCGACGTCTCTAACATGAGCCACTTGGCGAGGCAGAGACTGCTGGGCCGATCGTGGAGCGTGCCGGTCATCCGCCACCTCT-3’) were injected into fertilized single-cell stage embryos. Two-cell stage embryos were then transplanted into pseudo-pregnant female mice. Viable offspring were genotyped by next-generation sequencing. The *Dnmt3a^R878H/+^* mice were continually backcrossed with C57BL/6 mice to prevent the occurrence of epigenetic drift. Generation of the Puma-tdTomato reporter mouse has been previously described [38].

### Peripheral blood analysis

Peripheral blood (PB) was collected by retro-orbital or mandible venous puncture from live mice, or by heart puncture of deceased mice. PB was dispensed into EDTA-coated microvettes (Sarstedt #20.1288), and 200 µL of PB was then analysed using an ADVIA 2120 or 2120i (Siemens).

### Low coverage DNA sequencing for methylation analyses

Body tissues and select brain regions were harvested from 4- to 5-month-old *Dnmt3a^R878H/+^* mice or WT control littermates. DNA extraction was performed using a Zymo Quick-DNA™ Miniprep Plus Kit (Zymo #D4069). DNA quantification was performed using a NanoDrop, before samples were equalised to 10 ng/µL. Libraries were prepared according to the manufacturer’s instructions using the Oxford Nanopore Technologies Rapid Barcoding kit (SQK-RBK114-96). The libraries were then loaded onto a R10.4.1 PromethION flow cell (FLO-PRO114M) and sequenced at 0.01-0.05X coverage on a Nanopore sequencer, following the Skim-seq method of Faulk 2023 [20]. Base calling was performed with Dorado 0.7.0 (https://github.com/nanoporetech/dorado) using the super-accuracy model (dna_r10.4.1_e8.2_400bps_sup@v5.0.0) and the modified base call model 5mC_5hmC@v1. Base modifications calls were extracted for different sequence contexts (CG, CA, CAC) with modkit v0.3.1 (https://github.com/nanoporetech/modkit), setting the threshold for base modification calling to 0.8 (--filter-threshold C:0.8). Base modification calls for autosomes were aggregated to calculate the overall modification level for each sample. Samples with insufficient sequencing coverage, i.e. fewer than 70,000 measured cytosines per context (CG, CAC) equivalent to about 0.003X coverage, were discarded from the analysis.

### γ-radiation induced thymic lymphoma

Starting at the age of 4 weeks, *Dnmt3a^R878H^*^/+^ mice or WT control littermates were administered low dose (1.5 Gy) γ-irradiation, once a week for 4 weeks (total 4 doses). Trained technicians, blind to mouse genotype, identified and euthanised mice that had reached the predetermined ethical endpoint. Subsequently, thymic lymphoma was determined by abnormally increased thymus weight and/or abnormal CD4 and CD8 expression. Lymphoma-free survival was calculated in days from the last dose of γ-irradiation.

### Hematopoietic reconstitution and competitive bone marrow transplantation assays

Recipient mice on a C57BL/6-Ly5.1/Ly5.2 (CD45.1^+^CD45.2^+^) genetic background were lethally-irradiated with 2 doses of 5.5Gy, 2 h apart. They were then injected intra-venously (i.v.) with haematopoietic stem and progenitor cells derived from fetal liver cells from E13.5 embryos or whole bone marrow by intravenous tail injection. Recipient mice received a maximum of 2×10^6^ cells in a maximum volume of 200 uL of PBS intravenously. Following successful reconstitution, mice were monitored weedkly by animal technicians for signs of illness (as above).

Bone marrow was harvested from two femora of donor mice. The bone marrow was gently flushed and cells counted. Competitor bone marrow derived from *C57BL/6/Ly5.1J* (CD45.1^+^) mice was pooled together. The CD45.2^+^ bone marrow was harvested from either control WT mice or *Dnmt3a^R878H/+^* mice. The CD45.2^+^ cells were not pooled to maintain separate biological replicates. Competitor (CD45.1^+^) and test (CD45.2^+^) bone marrow cells were mixed in PBS.

For serial competitive bone marrow reconstitutions, recipient mice were left to age for 3 months following transplantation. At the end of the primary endpoint, bone marrow was harvested and cells counted. Bone marrow cells from each technical replicate were pooled in equal ratios to minimize technical variation. A total of 2×10^6^ bone marrow cells from each mixture were injected i.v. into each lethally γ-irradiated secondary recipient mouse. Blood, spleen, thymus and bone marrow were also collected for analysis by flow cytometry and histology.

### Flow cytometry and FACS

Single cell suspensions were generated for each haematopoietic organ of interest, before cells were stained with a cocktail of antibodies against cell surface markers with anti-FCR block (1:10, made in-house) in wash buffer (10% foetal bovine serum (SAFC) in PBS (Gibco)), protected from light on ice for 30 minutes. Immediately prior to analysis by flow cytometry or fluorescence-activated cell sorting (FACS), live cells were treated with a viability marker PI (2 μg/mL, Sigma-Aldrich), DAPI (1 μg/mL, BioLegend), Flourogold (1 μg/mL, AAT Bioquest), or TO-PRO-3 Stain (1 µM, ThermoFisher Scientific). Flow cytometry was performed on either an LSRII, Fortessa1, or Fortessa X20 (BD Biosciences) or Aurora (Cytek). Cell sorts were performed by staff at the WEHI FACs facility using a FACSAria III or FACSAria Fusion 9 (BD Biosciences). All data was analysed using FlowJo v10 (BD Biosciences). The cell types identified are defined in **Supplementary Table S1**, and all antibodies used are listed in **Supplementary Table S2**.

### RNA seq

4-week-old female *Dnmt3a^R878H/+^* mice or WT control littermates received a single low dose of γ-radiation (2 Gy) or were left untreated (baseline). 2 hours after γ-irradiation, mice were sacrificed, and their bone marrow used to generate single cell suspension, which were then stained with biotinylated antibodies against mature lineage surface markers. Magnetic selection using MagniSort Streptavidin Negative Selection Beads (Thermo Fisher) was used to remove mature leukocytes, as per the manufacturer’s protocol. The lineage-negative enriched bone marrow was then incubated with antibodies against c-KIT and SCA-1 to identify HSPCs, as well as antibodies against the mature lineage surface markers (as above) to detect any non-depleted mature lineage cells. Cells were finally stained with PI (2 μg/mL) and sorted (staining and sorting performed as above).

RNA from LSK cells was extracted using an miRNeasy kit (Qiagen #217084) with optional on-column DNase digestion (Qiagen #79254). Libraries were prepared using the Nextera XT DNA Library prep kit (Illumina) according to the manufacturer’s instructions.

Fourteen RNA-seq libraries were included in the analysis. Heatmaps were generated to show the expression levels of all genes in each of the selected pathways that were called enriched in the differentially expressed genes. Barcode plots depicting the enrichment levels of the pathways in our samples were generated using the barcode plot function in limma. The significance of the levels of enrichment was calculated by the roast function in the same package [PMID:20610611]. Gene tracks were created in Integrated Genomic Viewer open access package from the Broad Institute.

### Mito stress test

To determine the Oxygen Consumption rate (OCR) in cells, an XFe96 Extracellular Flux Analyzer was used. The assay was performed according to the manufacturer’s instructions in the Seahorse XF Cell Mito Stress Test Kit User Guide (103016-400, Agilent Technologies). 8.0×10^5^ B-cells or 6.0×10^5^ myeloid cells per well were plated on a Seahorse Plate for this assay. Baseline respiration was measured in XF RPMI medium (supplemented with 1 mM pyruvate, 2 mM glutamine, and 10 mM glucose). Oxygen consumption was further measured in differing metabolic conditions, following the addition of 3 μM Oligomycin, 2 μM FCCP and 2 μM Antimycin A/ 1μM Rotenone.

### Statistical Analysis and Figures

Prism software (Graphpad) was used to generate graphs, survival curves, and to analyze data unless otherwise indicated. Statistical methods used are referenced in the figure legends of relevant data. P value of <0.05 was deemed significant. If significance was tested and not indicated, results were non-significant. All data are presented as mean±SEM unless otherwise specified. Some figures were drawn using assets from Biorender.com.

## AUTHOR CONTRIBUTIONS

MJH and AW conceived the study. EML, AC designed and performed the experiments and data analysis. AJK, MP, LT generated the Dnmt3a^R878H^ mutant mice. EML, AC, HV, QG, JL, LW, SC performed experiments. EL, AT, AS provided the *Puma-TdTomato* reporter mice and helped with their use and interpretation of data from these experiments. AG, CLWS, WS, WL, QG, JL performed bioinformatic analysis. MJH, AS, AW provided expertise and supervision. The manuscript was drafted by EML, AC and MJH and edited by all co-authors. All authors reviewed the manuscript.

## DISCLOSURE OF CONFLICTS OF INTEREST

The authors declare no competing financial interests.

## ACKNOWLEDGEMENTS

This work was funded by NHMRC grant GNT1145728, a jointly funded PhD scholarship from Australian Rotary Health District 9650 and the Walter and Eliza Hall Institute, and the Alfred Hughes PhD Scholarship (WEHI). Grants for AS: NHMRC Research Fellowship GNT1116937; Grant-in-aid funding agreement between Cancer Council Victoria and Walter and Eliza Hall Institute of Medical Research, the estate of Anthony (Toni) Redstone OAM. Grants for MJH: NHMRC Investigator Grant 2017971, NHMRC Project Grants GNT1159658, GNT1186575, GNT1145728, and GNT1143105, and a Cancer Council Victoria Venture Grant. The generation of The *Dnmt3a^R878H/+^***^ ^**mice and *Puma-tdTomato* reporter mice used in this study was supported by Phenomics Australia and the Australian Government through the National Collaborative Research Infrastructure Strategy (NCRIS) program.

## SUPPLEMENTAL FIGURES

**<u>Figure S1:</u> Phenotyping of the *Dnmt3a^R878H/+^* mice**

**A)** Representative flow cytometry plots of *Dnmt3a^R878H/+^* and WT spleens. Leukocytes were gated by FSC-A and SSC-A, then cell doublets were gated out by FSC-H and FSC-A (not shown). Live cells were negative for dead cell markers PI, DAPI, or Fluorogold. Cells were defined based on cell type specific marker expression: CD4+ T cells = TCRβ+/CD4+/CD8-; CD8+ T cells = TCRβ+/CD4-/CD8+); transitional B cells = B220+/IgD_low_/IgM_high_; mature B cells = B220+/IgD+/IgM_mid_; granulocytes = TCRβ-/B220-/ Gr1+/Mac1+; and macrophages = TCRβ-/B220-/Gr1-/Mac1+. **B)** Percentages of haematopoietic cell subsets determined by flow cytometry in the blood and spleen. Data were statistically analysed using Student’s t-tests, *=p<0.05. **C-H)** Blood cell composition analyses from mice with irradiation induced thymic lymphoma: lymphocytes (C), neutrophils (D), monocytes (E), eosinophils (F), basophils (G), and large unstained cells (LUCs) (H). Data were statistically analysed using Student’s t tests, *=p<0.05. **I)** Flow cytometry plots of TCRβ and B220 expression in *Dnmt3a^R878H/+^*and WT spleen samples collected at the thymic lymphoma endpoint. Leukocytes were gated by FSC-A and SSC-A, then cell doublets were gated out by FSC-H and FSC-A (not shown). Live cells were negative for dead cell markers PI, DAPI, or Fluorogold. **J)** Flow cytometry plots of TCRβ and B220 expression in *Dnmt3a^R878H/+^* and WT thymus samples collected at the thymic lymphoma endpoint. Leukocytes were gated by FSC-A and SSC-A, then cell doublets were gated out by FSC-H and FSC-A (not shown). Live cells were negative for dead cell markers PI, DAPI, or Fluorogold. **K)** Graphical representations of the proportions of the cells described in J and K. Data were statistically analysed using a Mann-Whitney U test.

**<u>Figure S2:</u> Flow cytometry gating strategies, <u>Figure S3</u>: *ex vivo* γ-irradiation induced cell death analysis**

**A)** Flow cytometry gating strategy to define bone marrow HSPCs. Abbreviations: granulocyte myeloid progenitor (GMP), common myeloid progenitor (CMP), megakaryocyte erythroid progenitor (MEP), multipotent progenitor cell (MPP), long-term haematopoietic stem cell (LT-HSC), short-term haematopoietic stem cell (ST-HSC). **B)** Flow cytometric gating strategy to define HSPC compartments when assessing γ-irradiation-induced cell death. Live cells were determined via being negative for the dead cell marker Fluorogold.

**<u>Figure S3:</u> *Ex vivo* irradiation induced cell death analysis.**

**A-D)** Flow cytometric analyses of HSPC compartment was used to determine the cell numbers at each time point for myeloid-biased multipotent progenitor subset (MPP3) (A), short-term haematopoietic stem cells (ST-HSC) (B), long-term haematopoietic stem cells (LT-HSC) (C), and megakaryocyte-biased multipotent progenitor subset (MPP2) (D). For each condition, n=9 mice (3 mice per condition across 3 independent experiments). Error bars represent mean+/- SEM. Data were statistically analysed using 2-way ANOVAs and Tukey’s multiple comparisons test.

**<u>Figure S4:</u> RNAseq of untreated and γ-irradiated *Dnmt3a^R878H/+^* and WT LSK cells**

**A)** Schematic depicting the experimental workflow whereby LSK cells were isolated via FACs from the bone marrow, and their RNA extracted for RNAseq. **B)** MD plot depicting gene expression differences between non-irradiated *Dnmt3a^R878H/+^* and WT LSK cells. For the *Dnmt3a^R878H/+^* LSK cells, red and blue indicate significantly upregulated or downregulated genes, respectively, relative to the WT control. **C)** Heatmap depicting differentially expressed genes between non-irradiated *Dnmt3a^R878H/+^*and WT LSK cells. **D)** Gene ontology analysis reveals several downregulated pathways in untreated *Dnmt3a^R878H/+^* LSK cells. **E, F),** Barcode plots showing enrichment of genes in the p53 signalling pathway (KEGG pathway mmu04115) in non-irradiated WT LSK cells vs γ-irradiated WT LSK cells (E) and non-irradiated *Dnmt3a^R878H/+^* LSK cells vs γ-irradiated *Dnmt3a^R878H/+^* LSK cells (F).

**<u>Figure S5:</u> Oxygen consumption rates of bone marrow-derived *Dnmt3a^R878H/+^* and WT cells.**

**A)** Oxygen consumption rate (OCR) profiles of bone marrow-derived myeloid cells from *Dnmt3a^R878H/+^* (n=7) or WT (n=5) mice. **B)** OCR profile of bone marrow-derived B cells, *Dnmt3a^R878H/+^* (n=7) or WT (n=5) mice. **C)** Basal respiration rates for bone marrow-derived myeloid and B cells from *Dnmt3a^R878H/+^* (n=7) or WT (n=5) mice. **D)** Maximal respiration rates for bone marrow-derived myeloid and B cells from *Dnmt3a^R878H/+^* (n=7) or WT (n=5) mice.

**<u>Figure S6:</u> *Puma-tdTomato* reporter induction in thymocytes following y-irradiation**

**A)** Graphical representation *Puma-tdTomato* expression in the thymi of *Dnmt3a^R878H/+^* (n=5) and WT (n=5) mice, either untreated or after γ-irradiation. Data were statistically analysed using 2-way ANOVA with Šídák’s multiple comparisons test.

**Supplementary Table S1:** Defining cell markers for cell subsets.

| Cell Type | Cell Markers |
| --- | --- |
| LT-HSC | Lin-Sca1+cKIT+CD135/Flk2-CD34-;<br>Lin-Sca1+cKIT+CD135/Flk2-CD150+CD48- |
| ST-HSC | Lin-Sca1+cKIT+CD135/Flk2+CD34+;<br>Lin-Sca1+cKIT+CD135/Flk2-CD150-CD48- |
| MPP | Lin-Sca1+cKIT+CD135/Flk2+CD34+ |
| MPP2 | Lin-Sca1+cKIT+CD135/Flk2-CD150+CD48+ |
| MPP3 | Lin-Sca1+cKIT+CD135/Flk2-CD150-CD48+ |
| LSK | Lin-Sca1+cKIT+ |
| CLP | Lin-Sca1midKITmid |
| GMP | Lin-Sca1-cKIT+CD16/32+CD34+ |
| CMP | Lin-Sca1-cKIT+CD16/32-CD34+ |
| MEP | Lin-Sca1-cKIT+CD16/32-CD34- |
| Granulocytes | TCRβ-B220-Gr1+Mac1+ |
| Macrophages | TCRβ-B220-Gr1-Mac1+ |
| Pro-pre B | TCRβ-B220midIgM- |
| Immature B | TCRβ-B220midIgMmid |
| Transitional B | TCRβ-B220+IgM+ |
| Mature B | TCRβ-B220+IgM- |
| DN | CD4-CD8- |
| DN1 | CD4-CD8-CD25-CD44+ |
| DN2 | CD4-CD8-CD25+CD44+ |
| DN3 | CD4-CD8-CD25+CD44- |
| DN4 | CD4-CD8-CD25-CD44- |
| DP | CD4+CD8+ |
| CD4+ | CD4+CD8- |
| CD8+ | CD4-CD8+ |
| Lineage | CD4, CD8, CD11b/Mac1, Ter119, Ly6, B220, CD2, CD3, CD19, F4/80, NK1.1, Gr1 |

**Supplementary Table S2:** Antibody details.

| **Antibody** | **Fluorochrome** | **Clone** | **Source** |
| --- | --- | --- | --- |
| CD4 | PerCPCy5.5 | GK1.5-7 | eBioscience |
| CD4 | FITC | H129.19 | BioLegend |
| CD4 | A700 | GK1.5 | BioLegend |
| CD8 | BV650 | 53-6.7 | eBioscience |
| CD8 | A700 | 53-6.7 | BioLegend |
| TCRβ | PE-Cy7 | H57.597 | eBioscience |
| B220 | BV605 | RA3-6B2 | BioLegend |
| B220 | A700 | RA3-6B2 | BioLegend |
| Mac1/CD11b | APC-Cy7 | M1/70 | BioLegend |
| Gr1 | FITC | RB6-8C5 | BioLegend |
| Gr1 | A700 | RB6-8C5 | BioLegend |
| cKIT/CD117 | BV711 | 2B8 | BioLegend |
| cKIT/CD117 | APC | 2B8 | BioLegend |
| Sca1 | A594 | E13-161.7 | BioLegend |
| Sca1 | PE-Cy7 | D7 | BioLegend |
| CD19 | FITC | ID3 | eBioscience |
| Ter119 | PE | TER119 | eBioscience |
| Ter119 | A700 | TER119 | eBioscience |
| IgD | BV510 | 11.26c.2a | BioLegend |
| IgD | FITC | 11.26V | eBioscience |
| NK1.1 | A700 | PK136 | eBioscience |
| CD44 | FITC | IM781 | eBioscience |
| CD25 | PE | PC61 | BioLegend |
| Ly6G | A700 | 1A8 | eBioscience |
| CD2 | A700 | RM2-5 | eBioscience |
| CD3 | PE | 14-2C11 | eBioscience |
| CD3 | A700 | 17A2 | BioLegend |
| CD16/32 | PerCPCy5.5 | 242 | eBioscience |
| IgM | APC | MM-30 | BioLegend |
| IgM | PE-Cy5 | RMM-1 | BioLegend |
| CD34 | FITC | RAM34 | eBioscience |
| CD135 | PE | A2F10 | BioLegend |
| CD48 | FITC | HM48-1 | BioLegend |
| CD150 | BV421 | TC15-12F12.2 | BioLegend |
| CD45.1 | BV421 | A20 | eBioscience |
| CD45.2 | PE | 104 | eBioscience |
| F4/80 | A700 | BM8 | BioLegend |

## Notes

### Competing Interest Statement

The authors have declared no competing interest.

## REFERENCES

1. Fong, C.Y., J. Morison, and M.A. Dawson, Epigenetics in the hematologic malignancies. Haematologica, 2014. 99(12): p. 1772–83.

2. Venney, D., A. Mohd-Sarip, and K.I. Mills, The Impact of Epigenetic Modifications in Myeloid Malignancies. International Journal of Molecular Sciences, 2021. 22(9): p. 5013.

3. Yang, L., R. Rau, and M.A. Goodell, DNMT3A in haematological malignancies. Nature Reviews Cancer, 2015. 15(3): p. 152–165.

4. Xu, J., et al., DNMT3A Arg882 mutation drives chronic myelomonocytic leukemia through disturbing gene expression/DNA methylation in hematopoietic cells. Proceedings of the National Academy of Sciences, 2014. 111(7): p. 2620–2625.

5. Lu, R., et al., Epigenetic Perturbations by Arg882-Mutated DNMT3A Potentiate Aberrant Stem Cell Gene-Expression Program and Acute Leukemia Development. Cancer Cell, 2016. 30(1): p. 92–107.

6. Scheller, M., et al., Hotspot DNMT3A mutations in clonal hematopoiesis and acute myeloid leukemia sensitize cells to azacytidine via viral mimicry response. Nature Cancer, 2021. 2(5): p. 527–544.

7. Challen, G.A., et al., Dnmt3a is essential for hematopoietic stem cell differentiation. Nat Genet, 2011. 44(1): p. 23–31.

8. Jeong, M., et al., Loss of Dnmt3a Immortalizes Hematopoietic Stem Cells In Vivo. Cell Reports, 2018. 23(1): p. 1–10.

9. Challen, Grant A., et al., Dnmt3a and Dnmt3b Have Overlapping and Distinct Functions in Hematopoietic Stem Cells. Cell Stem Cell, 2014. 15(3): p. 350–364.

10. Lu, R., et al., A Model System for Studying the DNMT3A Hotspot Mutation (DNMT3AR882) Demonstrates a Causal Relationship between Its Dominant-Negative Effect and LeukemogenesisDominant-Negative Effect of DNMT3A R882 Mutation in Leukemia. Cancer research, 2019. 79(14): p. 3583–3594.

11. Ley, T.J., et al., DNMT3A Mutations in Acute Myeloid Leukemia. New England Journal of Medicine, 2010. 363(25): p. 2424–2433.

12. Guryanova, O.A., et al., DNMT3A mutations promote anthracycline resistance in acute myeloid leukemia via impaired nucleosome remodeling. Nature Medicine, 2016. 22(12): p. 1488–1495.

13. Dai, Y.-J., et al., Conditional knockin of Dnmt3a R878H initiates acute myeloid leukemia with mTOR pathway involvement. Proceedings of the National Academy of Sciences, 2017. 114(20): p. 5237–5242.

14. Liao, M., et al., Aging-elevated inflammation promotes DNMT3A R878H-driven clonal hematopoiesis. Acta Pharmaceutica Sinica B, 2022. 12(2): p. 678–691.

15. Wang, H., et al., Clonal hematopoiesis driven by mutated DNMT3A promotes inflammatory bone loss. Cell, 2024.

16. Smith, A.M., et al., Functional and epigenetic phenotypes of humans and mice with DNMT3A Overgrowth Syndrome. Nature Communications, 2021. 12(1): p. 4549.

17. Tuval, A., et al., Pseudo-mutant P53 is a unique phenotype of DNMT3A -mutated pre-leukemia, in Haematologica. 2022, Ferrata Storti Foundation: Italy. p. 2548–2561.

18. Tuval, A., et al., Pseudo-mutant P53 is a unique phenotype of DNMT3A-mutated pre-leukemia. Haematologica, 2022. 107(11): p. 2548–2561.

19. Wang, Y.A., et al., DNA methyltransferase-3a interacts with p53 and represses p53-mediated gene expression. Cancer Biol Ther, 2005. 4(10): p. 1138–43.

20. Faulk, C., Genome Skimming with Nanopore Sequencing Precisely Determines Global and Transposon DNA Methylation in Vertebrates. Genome Research, 2023(33): p. 948–956.

21. Moore, L.D., T. Le, and G. Fan, DNA Methylation and Its Basic Function. Neuropsychopharmacology, 2013. 38(1): p. 23–38.

22. He, B., et al., Tissue-specific 5-hydroxymethylcytosine landscape of the human genome. Nature Communications, 2021. 12(1): p. 4249.

23. de Mendoza, A., et al., The emergence of the brain non-CpG methylation system in vertebrates. Nat Ecol Evol, 2021. 5(3): p. 369–378.

24. Tan, H.K., et al., DNMT3B shapes the mCA landscape and regulates mCG for promoter bivalency in human embryonic stem cells. Nucleic Acids Res, 2019. 47(14): p. 7460–7475.

25. Mao, S.Q., et al., Genome-wide DNA Methylation Signatures Are Determined by DNMT3A/B Sequence Preferences. Biochemistry, 2020. 59(27): p. 2541–2550.

26. Li, Y., et al., Rapid and Accurate Remethylation of Dnmt3a Deficient Hematopoietic Cells with Restoration of DNMT3A Activity. Blood, 2023. 142: p. 4124.

27. Beard, D.C., et al., Distinct disease mutations in DNMT3A result in a spectrum of behavioral, epigenetic, and transcriptional deficits. Cell Rep, 2023. 42(11): p. 113411.

28. Kinde, B., et al., Reading the unique DNA methylation landscape of the brain: Non-CpG methylation, hydroxymethylation, and MeCP2. Proc Natl Acad Sci U S A, 2015. 112(22): p. 6800–6.

29. Mayle, A., et al., Dnmt3a loss predisposes murine hematopoietic stem cells to malignant transformation. Blood, 2015. 125(4): p. 629–638.

30. Kaplan, H.S., THE ROLE OF RADIATION ON EXPERIMENTAL LEUKEMOGENESIS. Natl Cancer Inst Monogr, 1964. 14: p. 207–20.

31. Haney, S.L., et al., Dnmt3a Is a Haploinsufficient Tumor Suppressor in CD8+ Peripheral T Cell Lymphoma. PLOS Genetics, 2016. 12(9): p. e1006334.

32. Ladle, B.H., et al., De novo DNA methylation by DNA methyltransferase 3a controls early effector CD8+ T-cell fate decisions following activation. Proc Natl Acad Sci U S A, 2016. 113(38): p. 10631–6.

33. Venugopal, K., et al., Alterations to DNMT3A in Hematologic Malignancies. Cancer Res, 2021. 81(2): p. 254–263.

34. Guryanova, O.A., et al., Dnmt3a regulates myeloproliferation and liver-specific expansion of hematopoietic stem and progenitor cells. Leukemia, 2016. 30(5): p. 1133–42.

35. Loberg, M.A., et al., Sequentially inducible mouse models reveal that Npm1 mutation causes malignant transformation of Dnmt3a-mutant clonal hematopoiesis. Leukemia, 2019. 33(7): p. 1635–1649.

36. Michalak, E.M., et al., Puma and to a lesser extent Noxa are suppressors of Myc-induced lymphomagenesis. Cell Death Differ, 2009. 16(5): p. 684–96.

37. Pietras, E.M., et al., Functionally Distinct Subsets of Lineage-Biased Multipotent Progenitors Control Blood Production in Normal and Regenerative Conditions. Cell Stem Cell, 2015. 17(1): p. 35–46.

38. Lieschke, E., et al., Mouse models to investigate in situ cell fate decisions induced by TP53 and other facto. EMBO, in press, 2024.

39. Villunger, A., et al., p53- and drug-induced apoptotic responses mediated by BH3-only proteins puma and noxa. Science, 2003. 302(5647): p. 1036–8.

40. Zhang, X., et al., Dnmt3a loss and Idh2 neomorphic mutations mutually potentiate malignant hematopoiesis. Blood, 2020. 135(11): p. 845–856.

41. Yang, L., et al., DNMT3A R882 mutation is associated with elevated expression of MAFB and M4/M5 immunophenotype of acute myeloid leukemia blasts. Leukemia & Lymphoma, 2015. 56(10): p. 2914–2922.

42. Jawad, M., et al., DNMT3A R882 Mutations Confer Unique Clinicopathologic Features in MDS Including a High Risk of AML Transformation. Frontiers in Oncology, 2022. 12.

43. Bond, J., et al., DNMT3A mutation is associated with increased age and adverse outcome in adult T-cell acute lymphoblastic leukemia. Haematologica, 2019. 104(8): p. 1617–1625.

44. Kramer, A.C., et al., Dnmt3a regulates T-cell development and suppresses T-ALL transformation. Leukemia, 2017. 31(11): p. 2479–2490.

45. Meyer, S.E., et al., DNMT3A Haploinsufficiency Transforms FLT3ITD Myeloproliferative Disease into a Rapid, Spontaneous, and Fully Penetrant Acute Myeloid Leukemia. Cancer Discov, 2016. 6(5): p. 501–15.

46. Chang, Y.I., et al., Loss of Dnmt3a and endogenous Kras(G12D/+) cooperate to regulate hematopoietic stem and progenitor cell functions in leukemogenesis. Leukemia, 2015. 29(9): p. 1847–56.

47. Yang, L., et al., DNMT3A Loss Drives Enhancer Hypomethylation in FLT3-ITD-Associated Leukemias. Cancer Cell, 2016. 29(6): p. 922–934.

48. Kim, S.J., et al., A DNMT3A mutation common in AML exhibits dominant-negative effects in murine ES cells. Blood, 2013. 122(25): p. 4086–9.

49. Russler-Germain, D.A., et al., The R882H DNMT3A mutation associated with AML dominantly inhibits wild-type DNMT3A by blocking its ability to form active tetramers. Cancer Cell, 2014. 25(4): p. 442–54.

50. Emperle, M., et al., The DNMT3A R882H mutation does not cause dominant negative effects in purified mixed DNMT3A/R882H complexes. Sci Rep, 2018. 8(1): p. 13242.

51. Emperle, M., et al., Mutations of R882 change flanking sequence preferences of the DNA methyltransferase DNMT3A and cellular methylation patterns. Nucleic Acids Res, 2019. 47(21): p. 11355–11367.

52. Buscarlet, M., et al., DNMT3A and TET2 dominate clonal hematopoiesis and demonstrate benign phenotypes and different genetic predispositions. Blood, 2017. 130(6): p. 753–762.

53. Kueh, A.J., et al., An update on using CRISPR/Cas9 in the one-cell stage mouse embryo for generating complex mutant alleles. Cell Death Differ, 2017. 24(10): p. 1821–1822.

