## Supplementary Figures for "Transcriptomic alterations including p53 pathway dysregulation prime DNMT3A mutant cells for transformation"

#### Supplementary Figure 1

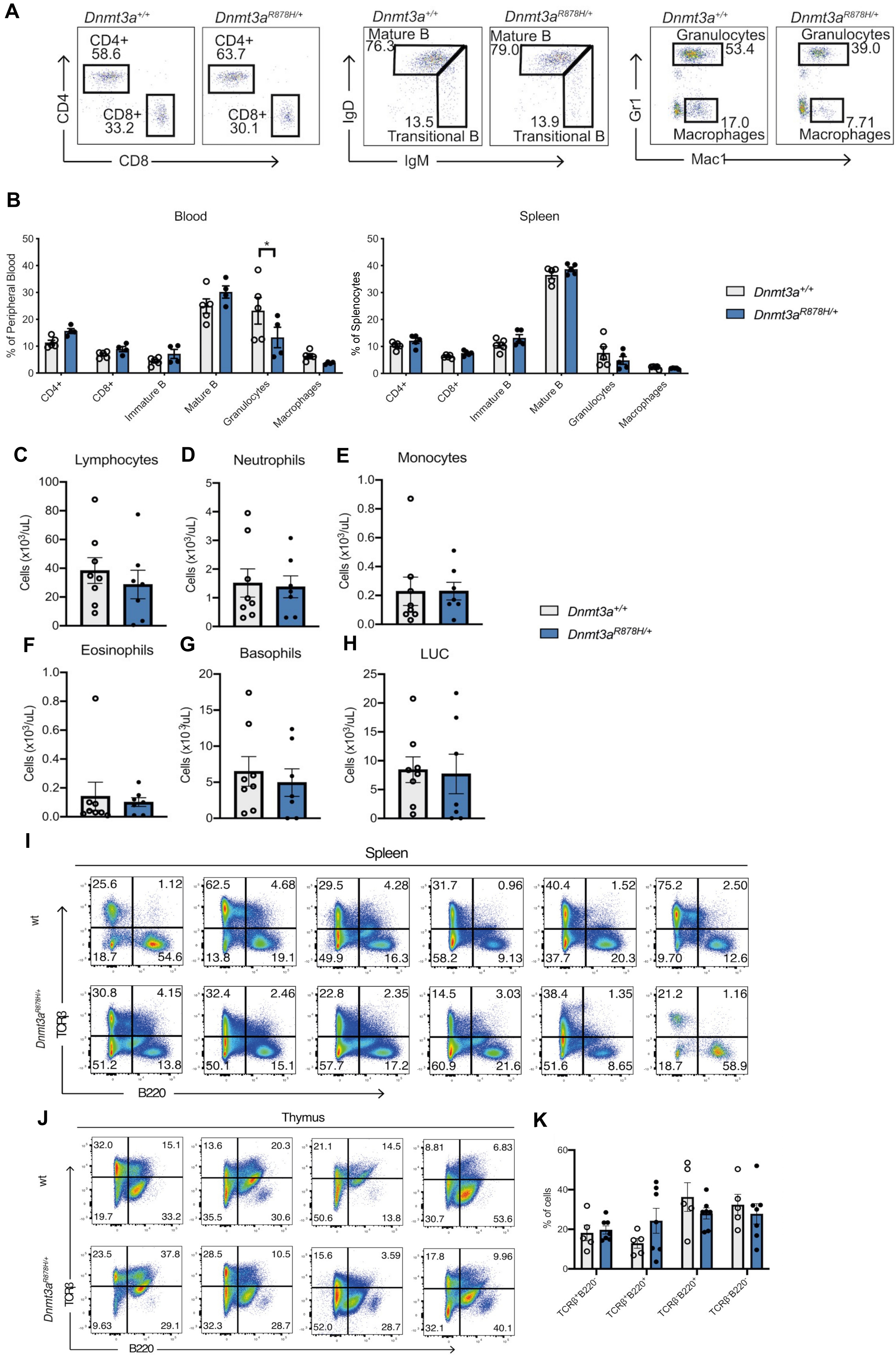

#### Supplementary Figure 2

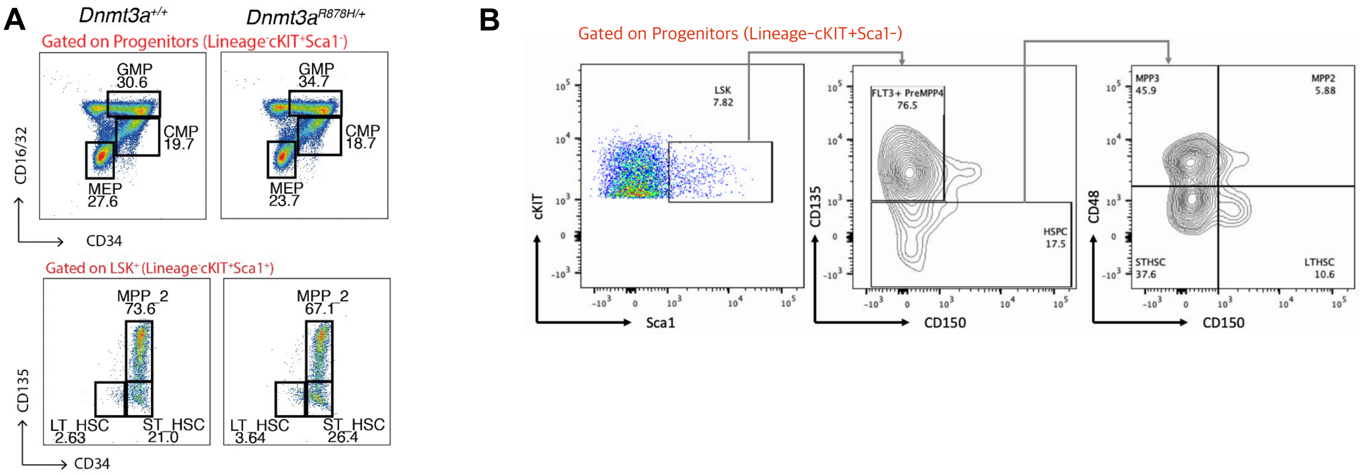

#### Supplementary Figure 3

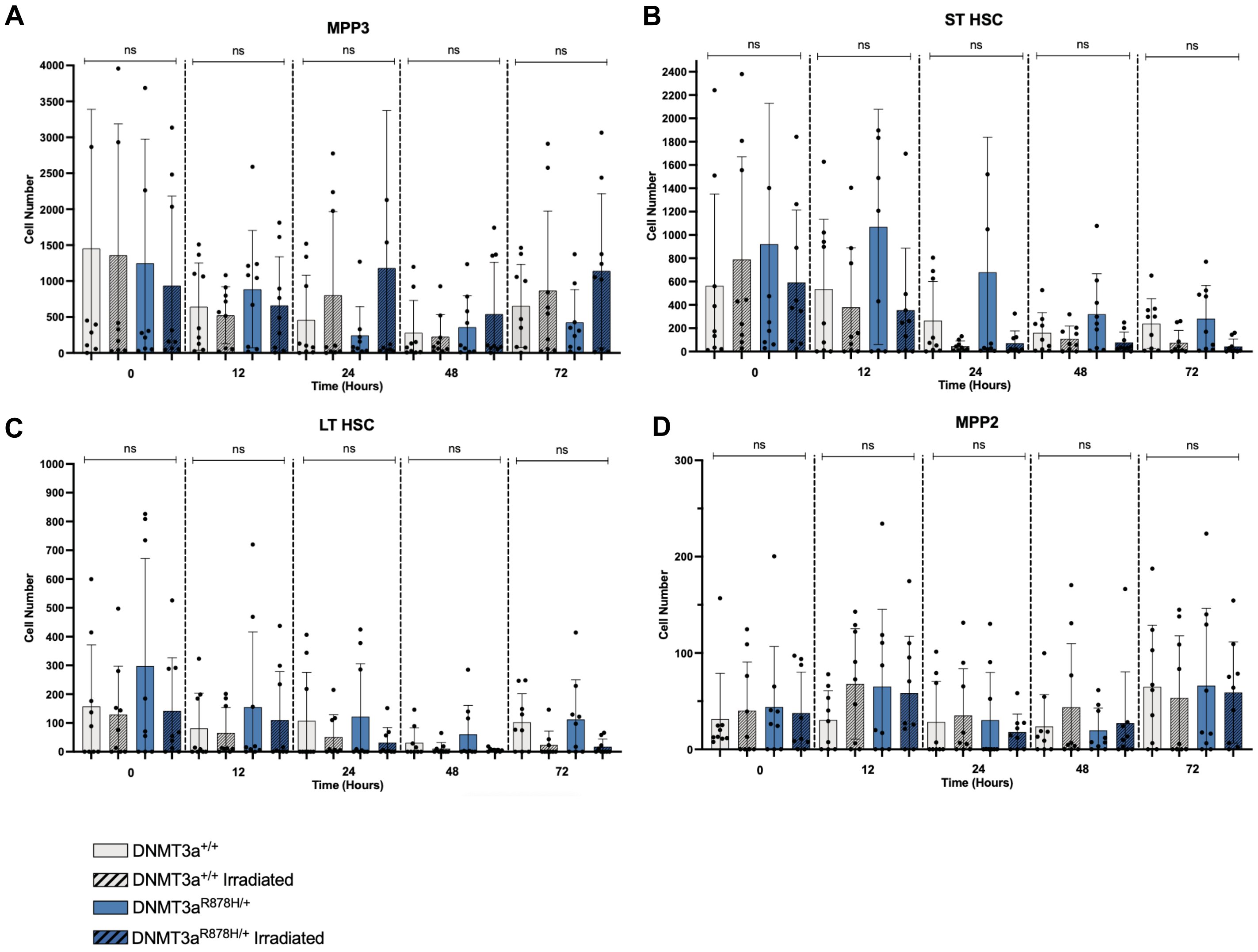

### Supplementary Figure 4

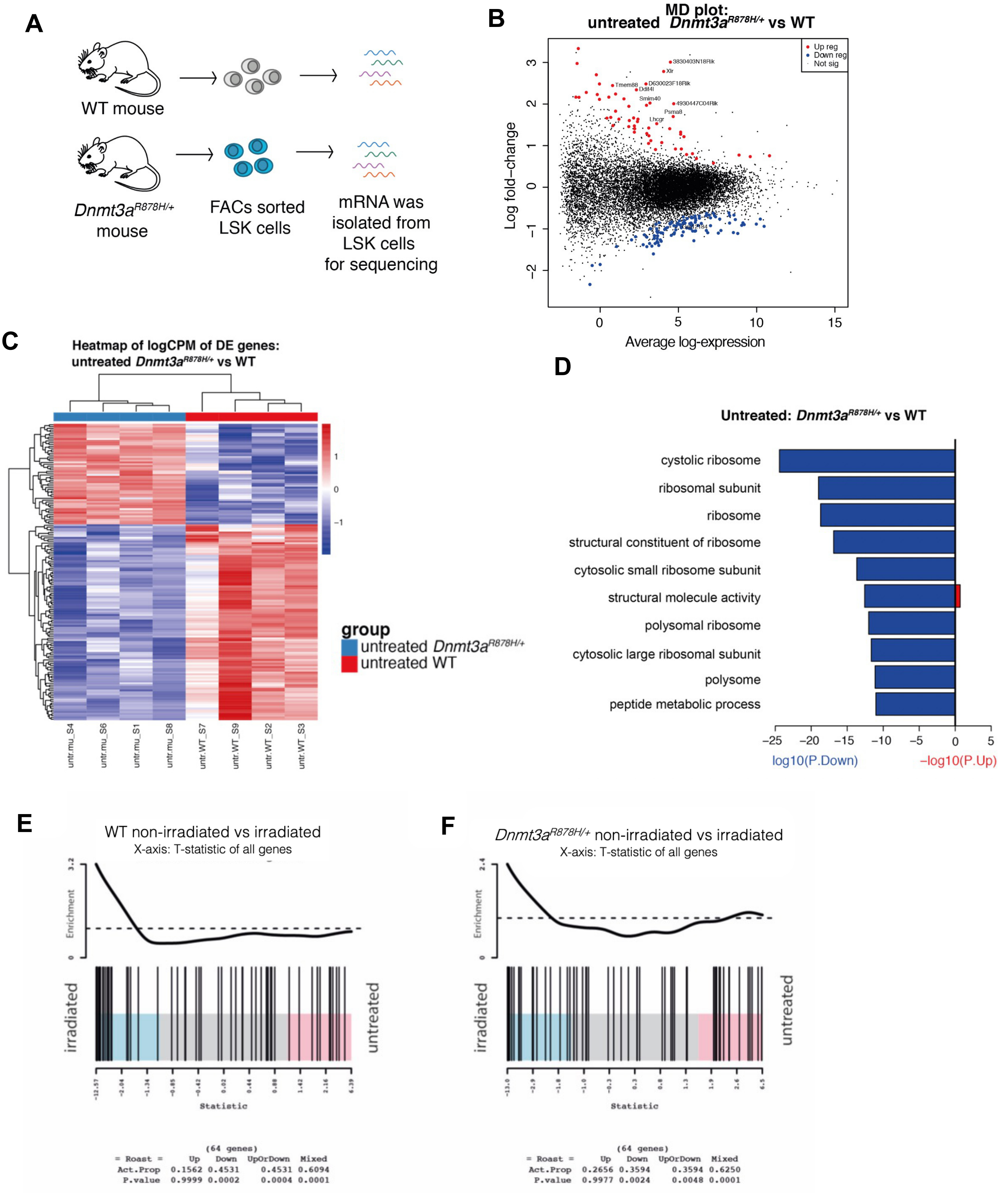

#### Supplementary Figure 5

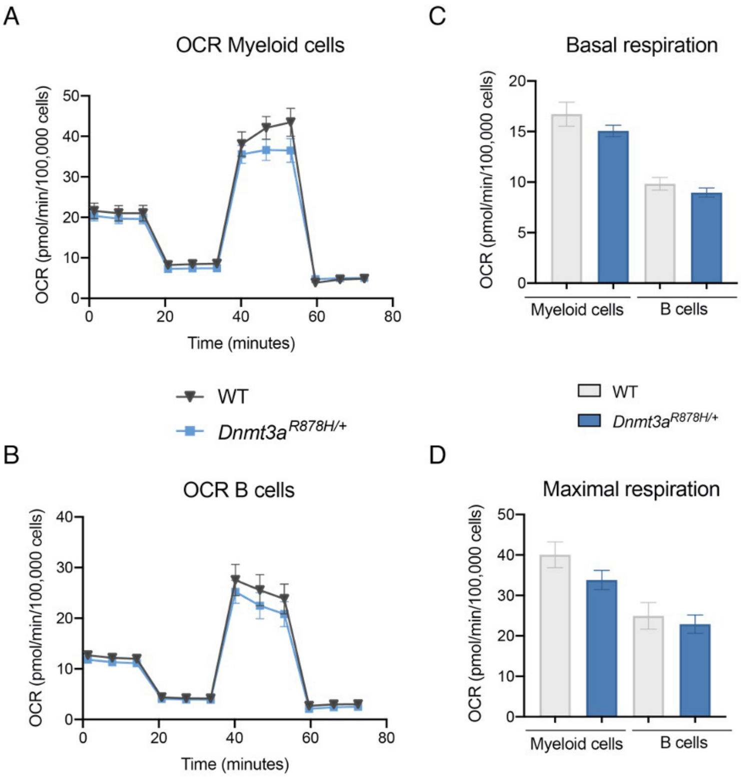

**Supplementary Figure 6**

A

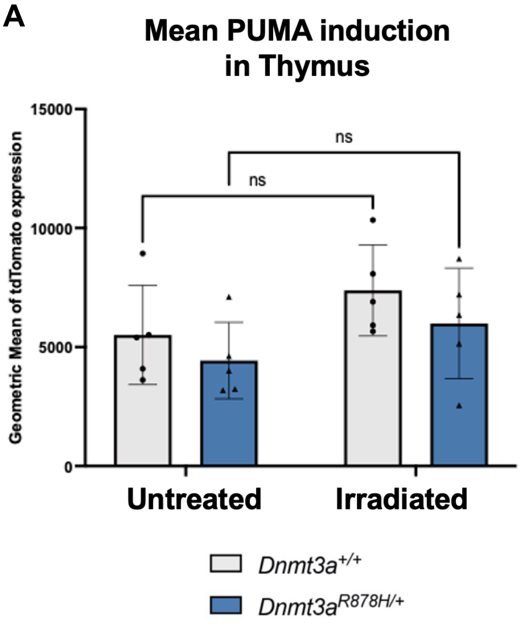
